# Movements and habitat selection of a marsupial carnivore in a modified landscape

**DOI:** 10.1101/2023.03.05.531208

**Authors:** Evie M. Jones, Amelia J. Koch, James M. Pay, Dydee F. Mann, Menna E. Jones, Rodrigo K. Hamede

**Affiliations:** University of Tasmania; Forest Practices Authority

## Abstract

Landscape modification is a major threat to carnivores worldwide, but modified landscapes can also provide important habitat for these species, as protected areas alone are insufficient. Understanding how carnivores use modified landscapes, such as production forests, can inform management strategies to improve the value of these landscapes to carnivores. Little is known about habitat selection by marsupial carnivores in production forests, where they occupy a similar ecological niche to their more well-studied eutherian counterparts. We used GPS tracking, Hidden Markov Models, and Manly’s selection ratios to identify the habitat selected in 3 behavioral states by the largest marsupial carnivore, the Tasmanian devil (*Sarcophilus harrisii*), in a timber plantation-dominated landscape. Behavioral states were approximated as: state 1, resting or feeding at a carcass; state 2, foraging; and state 3, travel. Devils did not show preferences for any of native forest, native grassland, and plantation in any behavioral state. Within plantations, devils preferred a plantation age of 4–7 years (selection ratio [*wi*] = 1.52). Devils preferred roads (state 1: *wi =* 2.71, state 2: *wi* = 2.48, state 3: *wi* = 2.97) and plantation edges (state 1: *wi =* 2.38, state 2: *wi* = 2.24, state 3: *wi* = 2.78) in all behavioral states, and moved faster on roads and edges than away from them. Together, our results indicate devils use road and edges for foraging (scavenging and hunting) and travel. No measured habitat variables influenced devil home range size. To support devils in plantation landscapes, we recommend maintaining a heterogeneous landscape of different plantation ages and native remnants and reducing the risk of vehicle collisions by minimizing forestry traffic at night. Tasmanian devils share similar adaptable traits to generalist eutherian carnivore species in their use of modified landscapes. Plantations can provide valuable habitat for this and other threatened predator species.

## Introduction

With the ongoing global loss of untouched forests (Curtis et al., 2018), there is increasing focus on finding strategies to preserve and enhance biodiversity in human-modified landscapes (Brockerhoff et al., 2008). Modified habitats such as production forests can make an important contribution to maintaining forest ecosystems, as protected areas alone are insufficient (Hayes and Ostrom, 2005). For instance, replacing native forests with timber plantations (henceforth ‘plantations’) negatively affects forest biodiversity (Kanowski et al., 2005; Wang et al., 2022), but plantations often contain higher biodiversity than other land uses such as agriculture, and can decrease harvesting pressure on native forest (Brockerhoff et al., 2008). Conservation strategies in production forests should be informed by ecological research to maximize effectiveness and cost-efficiency (Hartley, 2002).

Carnivores are especially sensitive to modified landscapes (Ripple et al., 2014) and knowledge of their habitat use within these landscapes can inform practices to sustain their populations. This is particularly important as carnivores have key ecological effects such as limiting populations of prey and smaller predators (Ritchie and Johnson, 2009), effects that are dampened in modified landscapes due to low densities of carnivores (Kuijper et al., 2016). Improving carnivore conservation in anthropogenic landscapes such as production forests can therefore benefit entire ecosystems in these landscapes. Carnivore species can differ markedly in both their habitat selection within plantations (Lantschner et al., 2012; Moreira-Arce et al., 2016) and their responses to forestry operations (Escudero-Páez et al., 2018). While the use of production forests by eutherian carnivores has been well studied (Ferreira et al., 2018), little is known about the use of these landscapes by marsupial carnivores, which occupy a similar ecological niche and face comparable threats from habitat loss (Jones & Barmuta, 2000; McGregor et al., 2014).

As well as displaying broad preferences among habitats, carnivores may also show finer scale differences in the habitats used for different behaviors, such as travel, foraging and denning (Houle et al., 2010). These behaviors can be investigated by looking at an animal’s movement paths – for instance, tortuous movements may indicate foraging (such as searching for prey or carrion) (Fuller and Harrison, 2010), while more directional movement may indicate travel (Andersen et al., 2017a). Understanding carnivore movements within these landscapes can help elucidate these behaviors and highlight important habitat characteristics (Fuller and Harrison, 2010). Animals often change their movements in response to altered forest structure (Moriarty et al., 2016). Habitat features such as windrows (linear piles of logging debris) (Sullivan and Sullivan, 2014) and corridors (MacDonald, 2003) can facilitate animal movement across production forest landscapes, and plantations can connect patches of native forest (Lyra-Jorge et al., 2008). Anthropogenic linear features in production forest landscapes, such as roads and edges between natural and modified vegetation, can be used by carnivores for travel and hunting (Bojarska et al., 2017; Gurarie et al., 2011) but also expose them to threats such as vehicle collisions (Jones, 2000). Investigating how carnivores move through production forestry landscapes, and the behaviors influencing habitat selection, can inform management strategies to promote important habitat features in the landscape.

With plantation forest cover in Australia increasing, fauna conservation is an important consideration in plantation management (Lindenmayer and Hobbs, 2004). This is particularly relevant in Tasmania where plantations comprise nearly 10% of total forest cover (Forest Practices Authority, 2017), and thus have significant potential to supplement native forest habitats and contribute to maintaining ecological function of the broader Tasmanian forest ecosystems if managed with conservation in mind.

Tasmania is home to the world’s largest extant marsupial carnivore, the Tasmanian devil. Devils are endangered due to a rare transmissible cancer, devil facial tumor disease (DFTD) (Hawkins et al., 2008), that has caused a steep population decline of over 80% since 1996 (Cunningham et al., 2021), leaving them vulnerable to other threats. Devils play multiple important roles in Tasmanian ecosystems: as morphologically specialized scavengers, they influence carrion persistence in the landscape (Cunningham et al., 2018), and as top predators, they help maintain trophic function and suppress feral cats *Felis catus* (Cunningham et al., 2020). Finding ways to protect devils in human-modified landscapes is a priority to maintain ecosystem function. Previous research investigated devil use of anthropogenic pastoral landscapes, finding that devils foraged along the pasture-forest interface and used pastures mainly for travel (Andersen et al., 2017a). Another study found devil abundance increases with plantation extent in production forest landscapes (Jones et al., 2023). Movements and habitat selection by devils in plantations have not been studied.

Our study provides the first analysis of marsupial carnivore habitat selection and movements within plantations. We used GPS tracking to explore the use of a plantation landscape by devils, with the intent to provide recommendations for plantation managers to sustain devil populations. We focused mainly on adult female devils, a key demographic to protect in critically diminished populations, particularly when they are raising young (April to January) (Bryant, 1988; Pemberton, 1990). We aimed to answer 4 questions:

1. At a plantation landscape scale, do devils select among different habitat types for different behaviors?
2. How do different habitats within plantation landscapes influence devil home range sizes?
3. At a plantation-specific scale, how do female devils select among plantation ages?
4. What features of plantations do devils use to move through the landscape?

This research complements similar studies on eutherian carnivores, enhancing our understanding of the use of production forests by carnivores and informing improved carnivore management in modified landscapes.

## Methods

### Study area

This study was undertaken in Surrey Hills, a eucalypt plantation landscape in northwest Tasmania (–41°S, 145°E) (Figure 1). The site is embedded in an intensively managed forestry region containing a mosaic of *Eucalyptus nitens* plantation stands of varying ages and remnant native forest and grassland patches. *E. nitens* is native to mainland Australia but not Tasmania. The site has an average annual rainfall of 2,163 mm and mean maximum temperature of 12.3°C (Bureau of Meteorology, 2022). The plantations have mostly been established since the mid-1970s on previously native vegetation land, and the remaining native vegetation primarily comprises highland *Poa* spp. grasslands, *Eucalyptus delegatensis* grassy forests and cool-temperate rainforest dominated by *Nothofagus cunninghamii* (Onfray, 2021). DFTD has been present in the landscape for about 6 years, causing steep population declines of devils (Cunningham et al., 2021).

**Figure 1:**
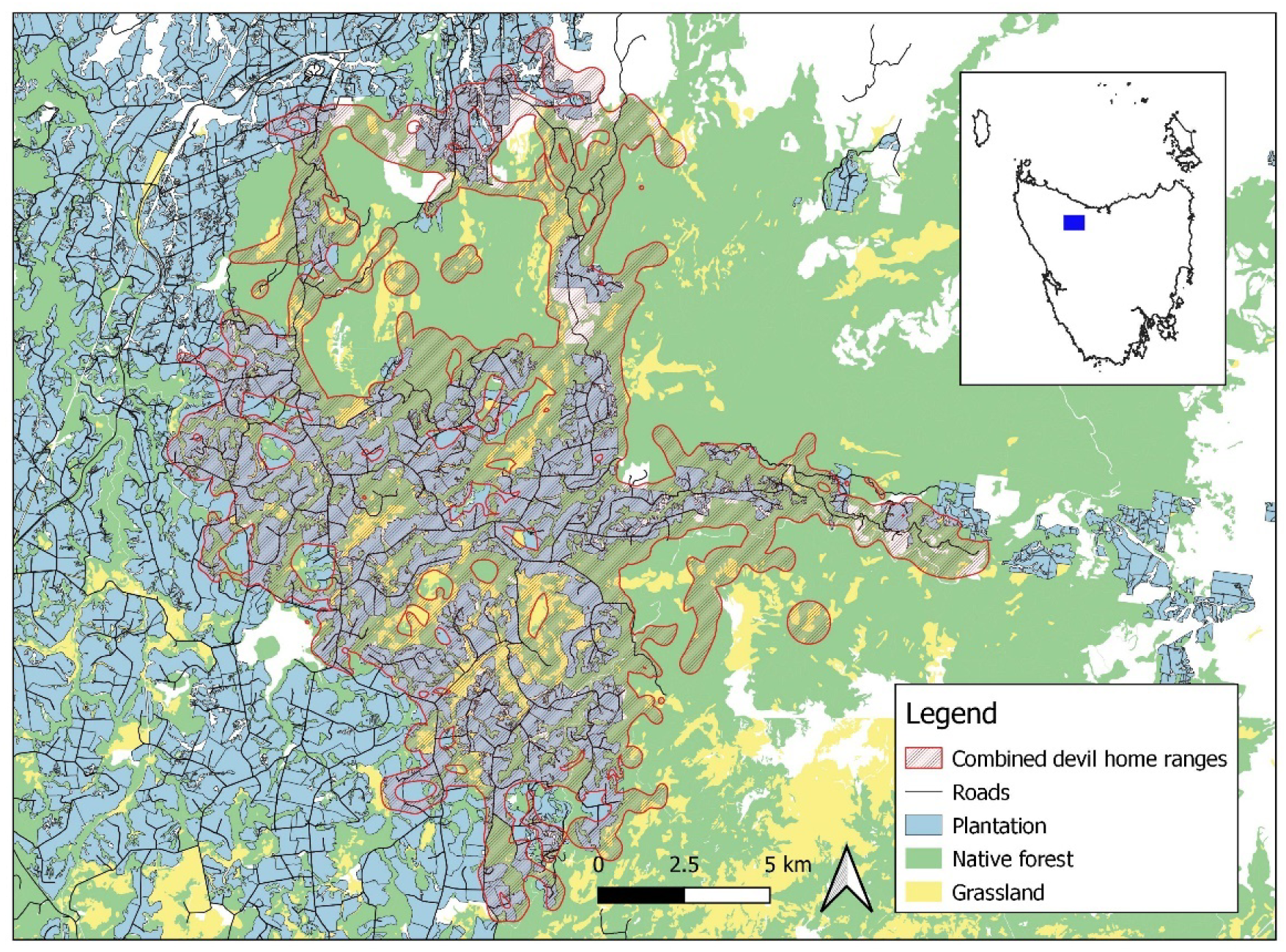
Study site within Tasmania (inset), overlaid with the combined home ranges of all devils in the study (95% utilization distribution from Biased Random Bridges).

### GPS telemetry

We captured adult devils (> 2 years old) in custom-built traps (made from 80 cm by 30 cm PVC pipe) baited with meat over 10-day trapping periods every month between April-October 2021. Traps were set within the plantation landscape in a mixture of plantation and native forest remnants. Monthly trapping allowed collared animals to be re-trapped to monitor collar fit and possible injuries. Upon capture, animals were weighed, sexed and checked for DFTD. We fitted adult devils with GPS radio-collars (Q4000ER GPS, Telemetry Solutions, Concord USA). We only collared animals that had been captured at least once before, indicating they were residents. Collars weighed 140 g, less than 3% of the body weight of collared individuals, and were fitted with corrosive bolts that degrade over time and eventually allow the collar to drop off (Thalmann, 2013). As devils are primarily nocturnal (Jones et al., 1997), collars were set to take 15 minute fixes every second night from approximately sunset until sunrise (17:30 to 07:00 April and August, 17:00 to 07:30 May to July, 18:00 to 06:00 September, 18:30 to 05:30 October) and were turned off during the day to conserve batteries. We set ‘GPS additional time’ to 30 seconds, which forces the GPS unit to keep searching for satellites even after it acquires a fix, improving fix accuracy. Collars were also equipped with a VHF unit to assist with relocation.

### GPS data screening

To account for errors in animal locations acquired by the GPS, fixes with less than 3 satellites, horizontal dilution of precision (HDOP) > 7 (as in Andersen et al., [2017a]), or unrealistic step lengths (> 4000 m) were removed from the analysis. Lower values of HDOP indicate higher location precision.

The successful fix rate of GPS radio-collars was low (23%), which can relate to factors such as species behavior (Mattisson et al., 2010), collar brand and/or habitat conditions (Di Orio et al., 2003; Hebblewhite et al., 2007), potentially confounding analysis. To account for potential habitat effects on fix success rate, we tested for bias in the success rate of GPS fixes due to vegetation type or plantation age. First, we left 10 collars stationary overnight in each of 3 vegetation types – native forest remnant, 1–5 year old plantation, and 10–15 year old plantation – and compared percentages of successful fixes among habitats. Second, we used straight-line interpolation to approximate the locations of failed fixes (‘approx’ function in R package ‘stats’) and compared the percentages of points in native forest, plantation and native grassland between real and interpolated points. Interpolated points were not used for analysis.

To investigate behavioral influences on GPS fix success rate, we plotted the ambient temperatures recorded by the collars and compared them between successful and failed fixes to evaluate how many failed GPS fixes could be attributed to animals resting in a den. Temperatures > 25°C appeared to indicate animals were in a den. Collar temperatures were unlikely to reach 25°C while animals were active since fixes were programmed only at night, and devils were collared over winter and spring in a relatively cold climate (annual mean max temp 12.3°C [Bureau of Meteorology, 2022]).

### Home ranges

We calculated devil home ranges using biased random bridges (BRBs) (Benhamou, 2011), which quantify an animal’s utilization distribution (UD) based on movement paths rather than just the density of individual fixes. We identified 95% UDs for each individual devil. Home ranges were also estimated using 100% minimum convex polygons (MCPs) (Mohr, 1947) for comparison with the broader literature. This is a commonly reported method of home range estimation, though it is prone to overestimating the areas actually used by animals (Burgman and Fox, 2003; Hemson et al., 2005). We did not use kernel density estimates (KDEs) (Worton, 1989), another widely used method, as the limited data and decentralized spread of GPS points for many individuals made this method unsuitable for our study. Analyses were carried out using the ‘adehabitatHR’ package in the R programming environment (Calenge, 2019).

We constructed maps of the study site using vegetation data from the TasVeg 3.0 layer (DPIPWE, 2017) and plantation features provided by the landowners, Forico Pty Ltd. Vegetation classes were grouped into 3 habitat types: plantation, native forest, and native grassland (including moorland and scrub). Plantation ages were grouped into 5 classes: < 1, 1–3, 4–7, 8–13 and 14+ years, age classes used by the forest industry due to different fuel loads. To examine if habitat variables influenced devil home range size (95% UDs), linear models were constructed using home range size as a response variable and proportions of different habitat types and plantation ages within each devil’s UD as predictors. These variables were included in separate models as proportions of different habitats/plantation ages would be correlated with one another, and plantation ages would be correlated with the percentage of plantation. Sex was also used as a predictor variable, both alone and in combination with other variables. As one male devil had a substantially larger home range than the other devils, we ran models with and without this individual to account for any skewed results. We selected final candidate models using Akaike’s Information Criterion values adjusted for small sample size (AICc) (Burnham and Anderson, 2002) according to the methods of Richards (2008), where models with ΔAICc > 6 were excluded, as were more complex models where simpler, nested models had smaller AICc values. To identify how much of the devils’ home ranges were affected by recent and active logging, and infer how this may influence their behavior, we calculated the percentage of each devil’s home range that was logged during the collaring period (April to October 2021) and in the 12 months preceding it.

### Behavioral states

To identify different behavioral states of female devils that may influence their habitat selection, we classified the GPS fixes for each animal into 3 different movement states using Hidden Markov Models (HMMs) (Patterson et al., 2008). This method classifies movement paths according to differences in turning angle and step length between successive locations, and assumes that these characteristics are driven by underlying behaviors. Movement states can therefore approximate behavioral states to some extent. Turning angle and step length are only calculated for successful fixes at consecutive 15 minute intervals, so successful fixes subsequent to failed fixes were excluded. We used past HMM results from devils to guide our choice of initial parameters for each state (Comte, 2019) (Table S1): state 1, stationary (e.g., resting and/or feeding at carcass); state 2, characterized by short step lengths and variable turning angles indicating tortuous movement (e.g., foraging); and state 3, characterized by long step lengths and relatively constant turning angles indicating directional movement (e.g., travel). Analyses were carried out using the ‘moveHMM’ package in R (Michelot et al., 2019).

Sequential GPS tracking data are often temporally autocorrelated which can violate assumptions for statistical inference (Boyce et al., 2010). To account for this, once behavioral states were assigned to all GPS fixes, we used variograms (function ‘variogram’ in R package ‘ctmm’ (Calabrese et al., 2016)) for individual devils in each state to find a time threshold at which variance stabilized and each location could be considered independent. GPS data variance stabilized and temporal autocorrelation decreased around three hours for state 1 and all combined states, and four hours for states 2 and 3. We subsampled the GPS data by these state-specific time thresholds (function ‘track_resample’ in R package ‘amt’; (Signer et al., 2019)) and used the subsampled data for habitat selection analysis. We did not have sufficient data to analyze selection of plantation ages by different behavioral states, so for plantation-specific habitat selection analyses, we repeated this process and subsampled the data by the time threshold for all combined states.

### Habitat selection and animal movements

For landscape-scale habitat selection analyses and for the plantation-specific analyses, we used the same ‘habitat type’ categories and ‘plantation age’ classes, respectively, as above. To investigate how animals may use plantation edges (edges between plantation and other habitat types, excluding roads) and roads for movement and foraging, we created three buffer zones around plantation edges and roads: < 20 m (on roads/edges, as in Andersen et al., [2017a]), 20–50 m (near roads/edges) and > 50 m (away from roads/edges). We calculated the percentage of each habitat type, plantation age class, and road/edge buffer in each devil’s 95% MCP (to reduce undue influence of outlying points), buffered by 5%, to represent available habitat. We used MCPs to define available habitat as we needed to measure all potentially available habitat rather than just the areas used by the animal. For each successful GPS point within the buffered MCPs we identified the habitat type, plantation age class (if applicable), and their distance to plantation edges and roads, split into the above buffers. These calculations were performed in QGIS 3.14^®^.

We used Manly’s selection ratios to investigate habitat selection by female devils in each HMM behavioral state, a method that tests for differences in used vs. available habitat (Manly et al., 2007). As there were insufficient male devils to examine sex differences in habitat selection, the three male devils were excluded from this analysis. We used a design III analysis, which recognizes that the available habitat differs among individuals (Manly et al., 2007). The selection ratio is the proportional use divided by proportional availability of each habitat attribute class, with a selection ratio (*wi*) > 1 indicating that a habitat type was used proportionally more than expected based on availability and < 1 proportionally less. The percentages of habitat attributes within buffered MCPs represented available habitat, while the locations of GPS points within the buffered MCPs represented used habitat. Univariate models were run for each behavioral state with the predictors of habitat type, plantation age, and distance to roads and plantation edges. Analyses were undertaken using the ‘adehabitatHS’ package in R (Calenge, 2011).

### Velocity

To support behavioral state analyses and help clarify devil movements in different habitats, we calculated the velocity of each devil at each GPS point using the difference in time and distance between two consecutive points using the ‘sp’ package in R (Pebesma et al., 2022). GPS points following a failed fix were ignored. The distance from each GPS point to the nearest road and plantation edge was calculated in QGIS 3.14^®^ and classified into ‘on road/edge’ (< 20 m), ‘near road/edge’ (20–50 m) and ‘away from road/edge’ (> 50 m). We tested for correlation between the values for plantation edges and roads using a polychoric correlation test (function ‘polychor’ in package ‘polycor’ in R [R Core Team, 2021]) and found low correlation (0.36), so these variables could be included in the same model. To investigate if devils move at different speeds in different habitats, we constructed linear mixed effects models using ‘velocity’ as a response variable and distance to road, distance to edge, habitat type, plantation age, and sex as predictors (function ‘lmer’ in package ‘lme4’ in R [R Core Team, 2021]). Interactions between habitat type/plantation age and distance to roads and edges were also included in models. Animal ID was included as a random effect. We ran 2 sets of models: ‘whole-of-landscape’ models, including all successful GPS fixes and all predictors except plantation age; and ‘plantation-only’ models, including only GPS fixes inside plantations, and all predictors except habitat type (to investigate the influence of plantation age). Final candidate models were selected using the same process as for the home range models (Burnham and Anderson, 2002), using AIC for whole-of-landscape models and AICc for plantation-only models.

## Results

We deployed GPS radio-collars on 15 adult devils (12 females, 3 males) between April and October 2021. One collar was unable to be retrieved at the end of the study, though some data were downloaded when the animal was recaptured during the study. A total of 7406 successful GPS fixes remained after removal of GPS errors, a success rate of 23% (successful vs programmed fixes). After accounting for temperatures recorded by the collar that indicated the animal was in a den (> 25 °C), the fix success rate increased to 52%, indicating much of the missing data was due to devils being in underground burrows where the collars could not detect satellites. Data were retrieved from between 24 and 164 days per animal (mean 77 ± 49 SD), with 152 to 1452 successful fixes per individual (494 ± 384) (Table S2). A total of 7066 points were used for analysis, following removal of points outside each animal’s buffered 95% MCP.

The stationary collar test had a fix success rate of 99% in plantation age 1–5 years, 97.2% in age 10–15 years, and 93.6% in native forest, similar to the difference in fix success rate among habitats in other studies (Andersen et al., 2017a). Comparison of the proportion of real vs. interpolated fixes that fell in each habitat category showed little difference between habitats (native forest 45.7% and 45.8% respectively, plantation 43.8% and 45.5%, grassland 10.4% and 8.8 %), indicating that any difference in fix success rate among habitats did not introduce a significant bias, though there was considerable variation among individuals (Table S3). The low successful fix rate in this study can therefore be attributed to animal behavior (denning) as well as other unknown factors. However, there is no evidence of significant habitat-induced bias, similar to the results of Mattisson et al. (2010), who investigated factors influencing fix success rate for GPS collars on wolverines (*Gulo gulo*).

The best performing HMM differentiated the 3 behavioral states as: state 1, representing little or no movement, with a mean step length of 16.37 ± 13.09 m (± SD) and a wide distribution of turning angles centered on 3.13 rad (concentration 0.39); state 2, with intermediate step lengths of mean 177.84 ± 148.25 m and a wide distribution of turning angles centered on 0.03 rad (concentration 0.52); and state 3, with longer steps of mean 527.03 ± 228.58 and a narrow distribution of turning angles centered on 0.03 rad (concentration 1.66), suggesting faster directional movement (Table S1, Figure S4). Subsampling the data by state-specific time thresholds to reduce spatiotemporal autocorrelation left 338 observations for state 1, 469 for state 2, 712 for state 3, and 859 for all combined states, which were used for habitat selection analysis.

### Habitat selection

At a landscape scale, behavior-specific habitat selection ratios for female devils showed that devils did not select among habitat types (plantation, native forest, and native grassland/scrub) in any behavioral state (state 1: *p =* 0.48, state 2: *p* = 0.69, state 3: *p* = 0.56) (Figure 2a). Female devils selected habitats non-randomly in all behavioral states regarding distance to roads (state 1: *p =* 0.01, state 2: *p* = 0.01, state 3: *p* < 0.001) and plantation edges (state 1: *p <* 0.001, state 2: *p* < 0.001, state 3: *p* < 0.001) (Figure 2c). Female devils used areas < 20 m from plantation edges (state 1: *wi =* 2.38 [95% CI 1.66–3.10], state 2: *wi* = 2.24 [1.63–2.85], state 3: *wi* = 2.78 [2.09–3.46]) and roads (state 1: *wi =* 2.71 [1.38–4.05], state 2: *wi* = 2.48 [1.59–3.37], state 3: *wi* = 2.97 [2.10–3.82]) more than twice as often as expected for all states. To a lesser extent, they selected for areas 20–50 m from plantation edges (*wi* = 1.47 [95% CI 1.11–1.82]) and roads (*wi* = 1.69 [1.19–2.20]) for state 2 only. They avoided areas > 50 m from edges (state 1: *wi =* 0.68 [95% CI 0.47–0.89], state 2: *wi* = 0.64 [0.49–0.79], state 3: *wi* = 0.62 [0.49–0.75]) and roads (state 1: *wi =* 0.87 [0.79–0.95], state 2: *wi* = 0.87 [0.81–0.92], state 3: *wi* = 0.89 [0.83–0.96]) in all states, particularly edges.

**Figure 2:**
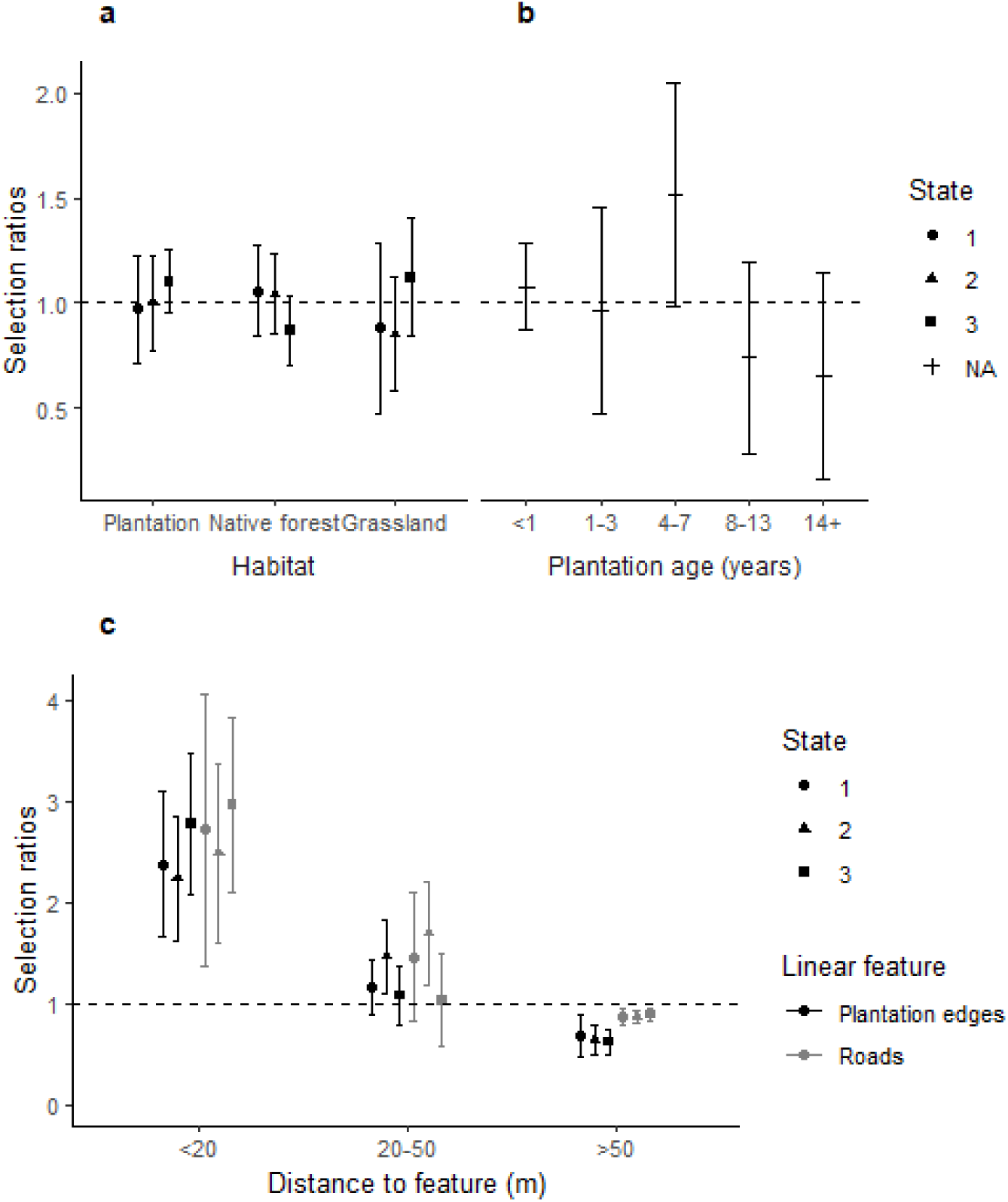
Habitat selection by adult female devils regarding habitat type (a), plantation age (b) and distance to roads/edges (c) in 3 behavioral states. Behavioral states were not considered for plantation ages due to limited data. Error bars show 95% CIs. A selection ratio of 1 indicates attributes were selected in proportion to availability; 0.5 indicates they were used half as often as expected; and 2 indicates they were used twice as often as expected, based on availability within buffered 95% MCPs.

Within plantations, habitat selection ratios for female devils indicated devils used plantations of different ages non-randomly (*p* < 0.001). Female devils selected for age 4–7 years (*wi* = 1.52 [95% CI 0.98–2.05]), and neither selected nor avoided young age classes < 4 years (*wi* = 0.96 [0.47–1.46]) and older age classes 8–13 (*wi* = 0.74 [0.28–1.20]) and 14+ years (*wi* = 0.65 [0.16–1.15]) (Figure 2b). All habitat selection results can be found in Table S5.

### Home ranges

Home range sizes (95% UDs from BRBs) were a mean of 1849 (416 SE) ha for female devils and 3343 (1388 SE) ha for males. 100% MCPs were a mean of 3481 (1475 SE) ha for females and 6326 (4401 SE) ha for males. UD and MCP estimations and plots for individual devils can be found in Table S2 and Figure S6. A single model remained in the final candidate set for predictors of devil home range size (95% UDs) and included only sex (Table 1). This remained the top model even when the male with the largest home range was removed from analysis, though the effect was weaker, with the null model in the final candidate list (ΔAIC = 2.85) (Table S7). Home ranges of males were on average 1494 (95% CI 566 to 2421) ha larger than females. The average recently logged area (< 1 year) in each devil’s home range was 3.46% (0–9.15%, Table S2). Male home range results should be treated with caution due to only 3 male devils in the study.

**Table 1:**
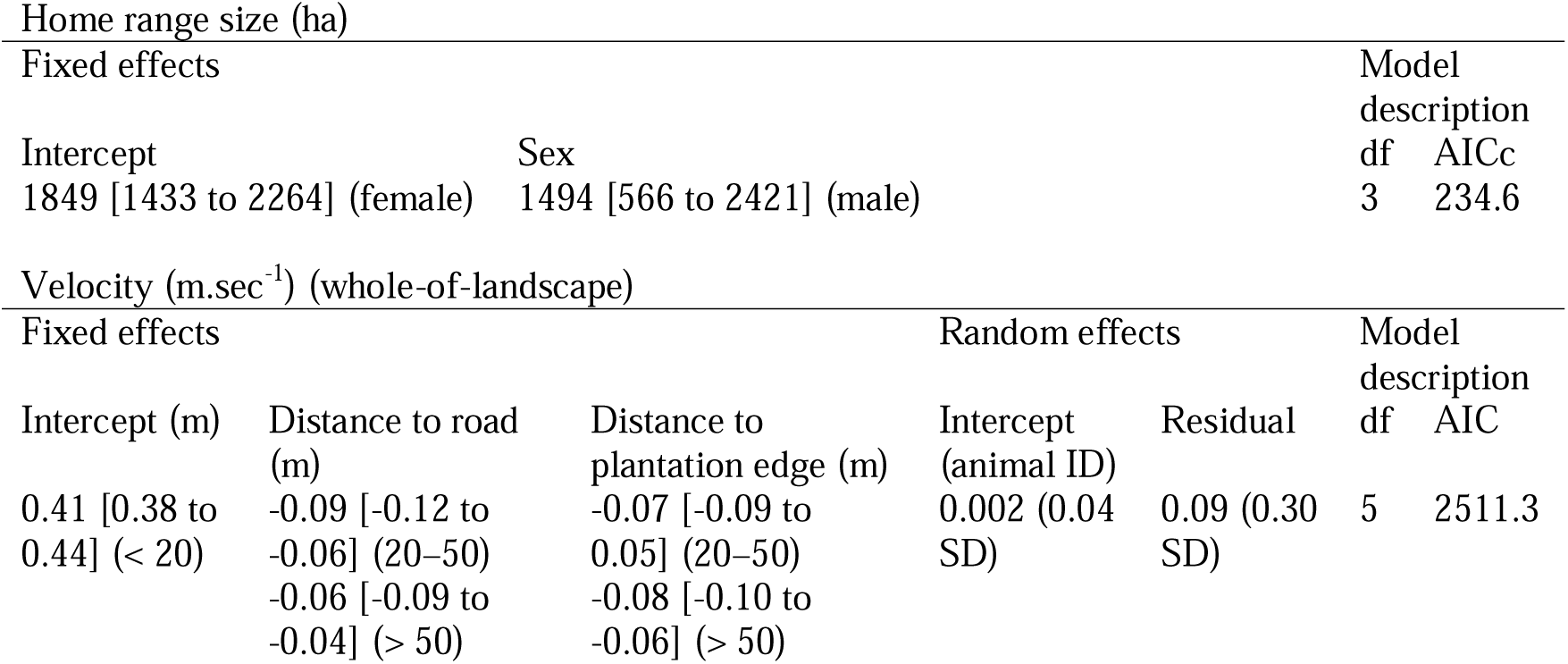
Final candidate models with parameter estimates for predicting devil home range size and velocity in plantations (95% CIs in square brackets []).

### Velocity

A single model remained in the final whole-of-landscape candidate set predicting devil velocity and this model included both distance to roads and plantation edges (Table 1). Devils moved faster on roads and edges (< 20 m) than away from them (Figure 3). In the plantation-only models, only distance to plantation edges remained in the final candidate list (Table S7).

**Figure 3:**
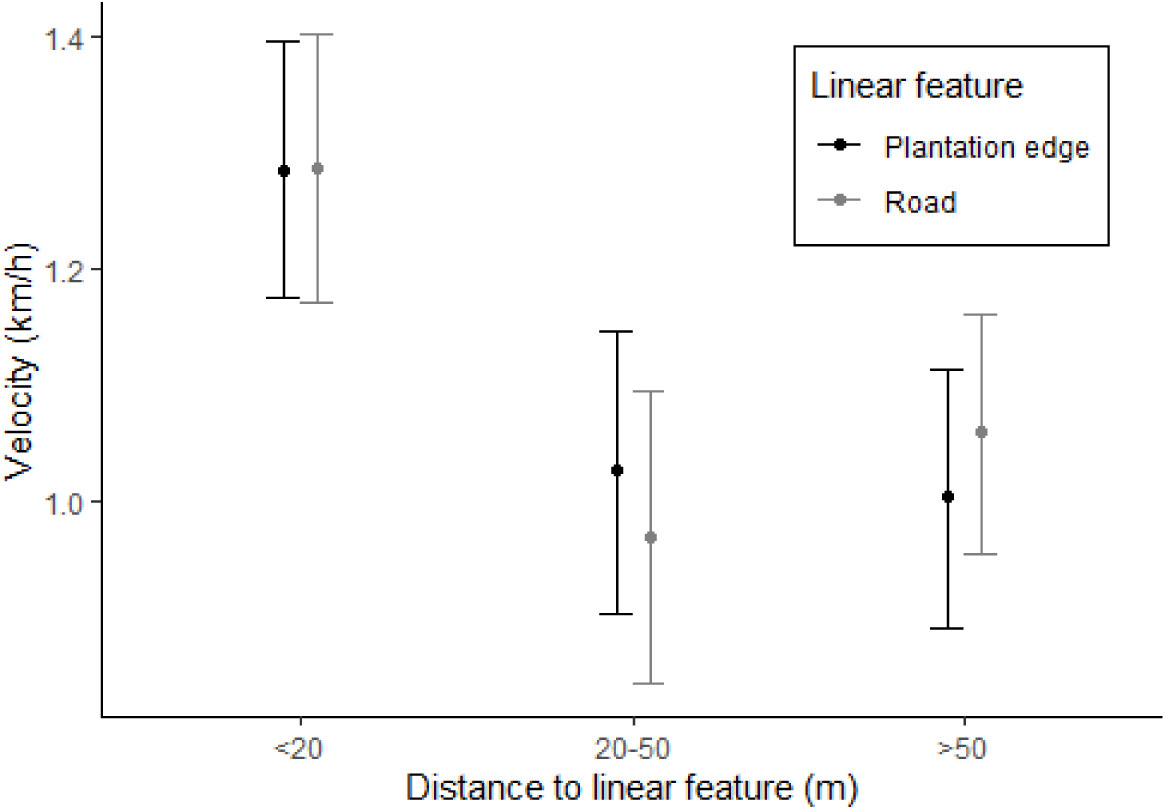
Predicted velocity of devils with distance to roads and plantation edges using whole-of-landscape data. Error bars represent 95% CIs.

As plantation-only models used a subset of the whole-of-landscape data and the results were similar, we consider whole-of-landscape results more relevant for interpretation.

## Discussion

Our results demonstrate the adaptability of devils to human-modified landscapes, highlighting the need to manage these landscapes as valuable devil habitat. Devils did not show preferences among different habitat types at a plantation landscape scale, though at a finer scale they showed selection for specific plantation ages, and they preferentially used plantation edges and roads to forage and to move through the landscape.

### Landscape-scale habitat selection and home ranges

At a landscape scale, female devils did not show selection among plantation, native forest remnants, and native grassland in any behavioral state, and no habitat attributes considered affected devil home range sizes. This demonstrates the adaptability of devils to human-modified landscapes, consistent with Jones et al. (2023)’s finding that devil abundance slightly increased with plantation extent in production forest landscapes. Devils are generalist carnivores (Andersen et al., 2017b), and generalist species usually respond more positively to human-modified landscapes than specialists (Dotta and Verdade, 2007). Other generalist carnivores such as raccoon dogs (Hwang et al., 2014) use plantations to a greater extent than native forest. Plantations likely provide abundant prey for devils. Medium-sized herbivores, wallabies and pademelons (Family Macropodidae; *Macropus rufogriseus* and *Thylogale billardierii*), and brushtail possums (*Trichosurus vulpecula*), preferred devil prey (Andersen et al., 2017b), are common enough in Tasmanian plantations that they are regularly culled to protect seedlings (Forest Practices Authority, 2017). Culling along with roadkill from forestry vehicles would provide abundant carrion for this specialized scavenger (Jones et al., 2003). There appear to be sufficient resources available in these landscapes for an opportunistic predator and facultative scavenger to exploit.

Notably, home ranges of female devils in this study were substantially larger than those in Andersen et al. (2020a) (100% MCPs: 2072 [277 SD] ha), which was conducted in an area that was free of DFTD at the time of the study and contains a mix of conservation and agricultural land. This is surprising as female devils decreased their home range sizes in another landscape that suffered significant population decline due to DFTD (Comte et al., 2020), likely due to less competition for resources. In the present study, we would therefore expect smaller home range sizes than in Andersen et al. (2020a) due to low devil density in our study landscape caused by high DFTD mortality (Cunningham et al., 2021). As home range size tends to increase with lower quality habitat (Kittle et al., 2015), this result may indicate that the plantation-dominated landscape in our study area was less ideal devil habitat than the more protected landscape in Andersen et al. (2020a). Perhaps protected landscapes provide more spatially and temporally constant and reliable resources (e.g., prey and den sites) than a production forest landscape that is always changing due to harvesting. For instance, black bears (*Ursus americanus*) were found to have larger home ranges in plantation dominated than in unmanaged forest landscapes (Jones & Pelton, 2003).

### Plantation-specific habitat selection

Within plantations, female devils selected for plantation aged 4–7 years. Younger plantations may contain a higher density of wallabies and brushtail possums which browse in plantations (Le Mar and Mcarthur, 2005). Some other carnivores, such as American marten (*Martes americana*) (Fuller and Harrison, 2005), similarly prefer younger post-logging native forest, likely due to higher prey availability. Additionally, near-surface vegetation cover in hardwood plantation, e.g., grass and shrubs, increase with plantation age in Tasmania (K. Parkins, pers. comm.), which may create a dense understory layer that restricts movement for devils in older plantations. While many carnivores prefer dense understory in plantations (Caryl et al., 2012; Escudero-Páez et al., 2018), this is often to avoid other predators or stalk prey. Devils are pounce-pursuit predators, capable of fast but short pursuit but dependent on some degree of ambush to get sufficiently close (Jones et al., 2003). They prefer forested rather than open environments and use forest edges to intercept herbivore prey as they move between daytime refuges and open areas to feed at night (Andersen et al., 2017b; Jones & Barmuta, 2000). The habitat structure offered by young plantations 4–7 years old may provide sufficient cover to conceal them from prey but not enough to significantly obstruct movement and potentially impede hunting and prey capture. Very young plantations may be too open for devils to be able to effectively ambush prey which could see them coming and move out of range.

It is interesting that female devils did not appear to avoid recently logged plantation (< 1 year), although this result was based on just 4 individuals. Considering female devils likewise did not avoid plantations 1–3 years old and actively selected for 4–7 years old, and that plantation forestry landscapes have high prey abundance, logging is unlikely to have a long-term negative effect on devil use of the landscape, as long as it is on a schedule that maintains plantation age heterogeneity in the landscape. Furthermore, logging removes habitat that is already apparently nonpreferred (coupes are logged at ∼12–15 years old in this landscape).

Other factors likely to influence the effect of logging on devils include the quality of remaining habitat in the surrounding area and home range (e.g., plantation ages and native forest remnants) (Lindenmayer and Hobbs, 2004); the timing of logging i.e., if logging occurs during the maternal denning season and could disrupt denning behavior; and the overall area and proportion of home range harvested (Leonard et al., 2008). We could not directly investigate how devils responded to active logging, however, the proportion of their individual home ranges that was logged during the study and in the preceding year was small (mean 3.5%, range 0–9%), so even short-term displacement due to logging may not have had a major effect on devils in this study. Extrapolation of these results to other years and landscapes should be treated with caution due to variations in logging area size and intensity. The lack of an apparent response to logging by devils is particularly surprising as the logging method in the landscape is clearfelling, which involves removing all trees from the harvest footprint and typically has a more negative effect on biodiversity than other logging methods (Chaudhary et al., 2016). Some carnivores can respond positively to clearfelling in plantations, perhaps due to increased prey availability (Eom et al., 2019).

### Devil movements in plantation landscapes

Devils preferentially used roads and plantation edges to move through the landscape in all behavioral states. This is consistent with Andersen et al. (2017b)’s discovery that devils use anthropogenic linear features for movement and foraging, and Andersen et al. (2020b)’s recording of devils scavenging along roads. Devils moved faster on roads and edges, indicating they use them for travel and perhaps hunting (Andersen et al., 2020b). Edges of plantation coupes are cleared of trees to allow vehicle access, potentially providing less obstructed movement paths for devils. Other carnivores such as wolves similarly use linear features such as roads and seismic lines for travel, allowing them to move further and faster (Dickie et al., 2017). Our results indicate devils also forage along roads and edges. As well as scavenging roadkill along roads (Andersen et al., 2020b, 2017a), devils may use plantation edges to intercept macropod prey moving between the denser shelter of native forest where they rest during the day to browse in more open plantations at night (Le Mar and Mcarthur, 2005), as was found for the forest-pasture interface in Andersen et al. (2017a). Other carnivores likewise select for edges in production forest landscapes (Houle et al., 2010; SimonsLLegaard et al., 2013), which can increase hunting success (Bojarska et al., 2017).

Female devils selected for areas close to (20–50 m), but not on, roads and edges for state 2 only. State 2 represents shorter, more tortuous movements which may indicate foraging, contrasted with longer, straighter step lengths for state 3 which suggests travel, and stationary behavior in state 1, i.e. resting or feeding at a carcass. Selection of areas 20–50 m from roads/edges in state 2 may be explained by devils searching for food near roads/edges but in cover to intercept prey, combined with a higher chance of detecting scents of prey or carcasses that are close to roads/edges while travelling along them.

### Management implications

As plantations appear to be valuable habitat for devils, we recommend implementing strategies to maintain habitat suitability for devils in plantation landscapes. Preserving a heterogeneous landscape with a mosaic of different plantation ages – along with patches of native vegetation remnants (Lindenmayer and Hobbs, 2004) – is likely to benefit devils due to their broad use of different habitat types and their preferences among plantation ages. Heterogeneous landscapes also contain a high density of habitat edges, which devils preferred in this study and in Andersen et al. (2017a). The use of plantation landscapes by devils exposes them to potential risks from human use of the landscape such as timber harvesting operations, vehicle collisions, and lead poisoning from consuming culled browser carcasses (Hivert et al., 2018). Our finding that devils preferentially use roads make them particularly vulnerable to vehicle collisions, a major cause of devil mortality (Jones, 2000). Logging in some coupes occurs during 24 hours in this landscape. Since measures targeted at changing driver behavior such as implementing speed limits and increasing driver awareness have limited effectiveness (Jones, 2000), the most effective way to mitigate this risk to devils may be to minimize forestry traffic at night. Our study suggests that the risk to devils from harvesting operations is likely to be minimal, as the age class devils preferred (4–7 years) is unlikely to be harvested. One caveat is that we were unable to investigate denning behavior of devils in this study, so the risk of damage to devil dens during logging operations – particularly maternal dens, which may harbor young devils unable to escape – remains unknown.

### Conclusions

Our study reinforces the high adaptability of devils to human-modified landscapes (Andersen et al., 2017a), similar to other generalist carnivores (Ferreira et al., 2018). Not only did devils show no selection between plantation and native habitat remnants, they took active advantage of modified habitat features – edges and roads – for foraging opportunities and expediated travel. Similar behavior has been observed before in devils (Andersen et al., 2020b, 2017a) as well as other carnivores (Dickie et al., 2017). Notably, this study only looked at habitat use of devils within a modified landscape, so future research would be necessary to identify the comparative value of modified and unmodified landscapes to devils.

Habitat selection by carnivores in production forests is highly variable among species, such as selection among harvested forest ages: some carnivores, like devils, prefer young-mid stage regrowth (Boisjoly et al., 2010), while others select for mature forest (Buskirk et al., 1996). Identifying species-specific habitat selection in human-modified landscapes is therefore important. While it is essential to recognize that carnivores as a group are particularly vulnerable to human impacts (Ripple et al., 2014), and that many carnivore species – particularly large and threatened carnivores – require unmodified habitats to thrive (Ferreira et al., 2018), it is also important to understand that many generalist carnivores are adaptable to modified landscapes, providing impetus to manage these landscapes as valuable habitat for these species. Production forests are an excellent example – they are often incorrectly viewed as of little value to conservation, exposing them to mismanagement and habitat degradation (Brockerhoff et al., 2008). Recognizing that production forests and other modified landscapes provide important habitat to devils and other carnivores is key to maintaining populations of these predators beyond protected areas.

## Notes

### Competing Interest Statement

The authors have declared no competing interest.

### Summary of Updates

Fixed abstract to match current version

