## Supplemental Information for "Movements and habitat selection of a marsupial carnivore in a modified landscape"

**Supporting information**

Appendix S1: Initial parameters for Hidden Markov Models for devil habitat selection.

|  | State 1 | State 2 | State 3 |
| --- | --- | --- | --- |
| Step mean (m) | 10 | 200 | 600 |
| SD | 10 | 200 | 300 |
| Zero-mass | 0.02 | 0.002 | 0.000001 |
| Angle mean (rad) | pi | pi | 0 |
| Concentration | 0.5 | 0.5 | 2 |

Appendix S2: Summary of data retrieved from GPS collared devils and their home range sizes. MCP = minimum convex polygon, BRB = biased random bridge.

| Name | Sex | Date collared | Days tracked | Successful fixes | Missing data (%) | MCP 100% (ha) | BRB 95% (ha) | BRB 50% (ha) | Logged (%) |
| --- | --- | --- | --- | --- | --- | --- | --- | --- | --- |
| Alannis | F | 18/04/2021 | 24 | 218 | 66.5 | 2260.49 | 1587.17 | 285.76 | 0.75 |
| Aphrodite | F | 8/05/2021 | 94 | 524 | 81.2 | 2034.70 | 1476.25 | 322.32 | 0.66 |
| Artemis | F | 21/04/2021 | 36 | 195 | 80.8 | 7392.98 | 2969.84 | 444.58 | 5.04 |
| Athena | F | 15/06/2021 | 35 | 207 | 79.4 | 2680.97 | 1590.47 | 288.93 | 0 |
| Doris | F | 16/04/2021 | 50 | 215 | 85.1 | 5050.49 | 1777.54 | 323.41 | 2.49 |
| Hella | F | 9/05/2021 | 20 | 152 | 69.8 | 3150.97 | 1628.51 | 227.25 | 2.63 |
| Hera* | F | 6/05/2021 | 156 | 1452 | 65.4 | 3605.17 | 2066.92 | 358.72 | 8.55 |
| Mike | M | 3/05/2021 | 71 | 499 | 75.1 | 11344.23 | 4901.29 | 577.59 | 3.44 |
| Naomi | F | 3/06/2021 | 79 | 340 | 85 | 2920.91 | 2016.61 | 269.01 | 0 |
| Pepper | F | 20/04/2021 | 61 | 330 | 80.2 | 2609.97 | 1660.3 | 264.13 | 2.02 |
| Prometheus | M | 3/05/2021 | 71 | 376 | 81.8 | 4509.26 | 2885.01 | 376.09 | 5.41 |
| Rocket | M | 3/05/2021 | 28 | 208 | 68 | 3123.16 | 2241.32 | 447.00 | 5.32 |
| Romanov | F | 16/06/2021 | 134 | 845 | 75.9 | 3416.45 | 1437.27 | 247.00 | 2.27 |
| Selene | F | 16/04/2021 | 131 | 753 | 78 | 2732.45 | 1977.05 | 393.08 | 4.10 |
| Wanda | F | 14/04/2021 | 164 | 1092 | 76.2 | 3920.89 | 1999.7 | 386.99 | 9.15 |

*collar removed 1/6/21-21/6/21

Appendix S3: Percentages of real and interpolated points for individual devils among different habitat types.

| Animal ID | Total real (%) | Total interpolated (%) | Plantation | | Native forest | | Grassland | |
| --- | --- | --- | --- | --- | --- | --- | --- | --- |
|  |  |  | Real (%) | Interpolated (%) | Real (%) | Interpolated (%) | Real (%) | Interpolated (%) |
| Alannis | 31.55 | 68.45 | 45.50 | 70.98 | 36.51 | 12.93 | 17.99 | 16.10 |
| Aphrodite | 17.21 | 82.79 | 41.69 | 46.73 | 42.13 | 32.58 | 16.19 | 20.69 |
| Artemis | 18.24 | 81.76 | 18.88 | 23.87 | 71.33 | 66.61 | 9.79 | 9.52 |
| Athena | 19.96 | 80.04 | 33.70 | 43.22 | 55.98 | 44.44 | 10.33 | 12.33 |
| Doris | 14.48 | 85.52 | 39.90 | 22.52 | 55.67 | 77.06 | 4.43 | 0.42 |
| Hella | 28.73 | 71.27 | 24.43 | 38.15 | 48.85 | 37.85 | 26.72 | 24.00 |
| Hera | 33.58 | 66.42 | 60.72 | 63.70 | 27.65 | 26.63 | 11.63 | 9.67 |
| Mike | 22.50 | 77.50 | 35.94 | 42.94 | 53.79 | 49.82 | 10.27 | 7.24 |
| Naomi | 14.86 | 85.14 | 60.29 | 59.18 | 33.46 | 38.64 | 6.25 | 2.18 |
| Pepper | 18.59 | 81.41 | 41.90 | 52.01 | 29.23 | 36.90 | 28.87 | 11.09 |
| Prometheus | 17.86 | 82.14 | 50.48 | 42.92 | 40.89 | 55.76 | 8.63 | 1.32 |
| Rocket | 32.09 | 67.91 | 42.63 | 54.98 | 53.16 | 42.79 | 4.21 | 2.24 |
| Romanov | 23.04 | 76.96 | 18.24 | 24.47 | 79.52 | 73.80 | 2.24 | 1.72 |
| Selene | 21.38 | 78.62 | 46.00 | 48.81 | 38.46 | 31.67 | 15.54 | 19.52 |
| Wanda | 21.81 | 78.19 | 44.11 | 44.49 | 53.35 | 52.48 | 2.54 | 3.03 |
| Mean | 22.39 | 77.61 | 40.29 | 45.26 | 48.00 | 45.33 | 11.71 | 9.40 |

**a**

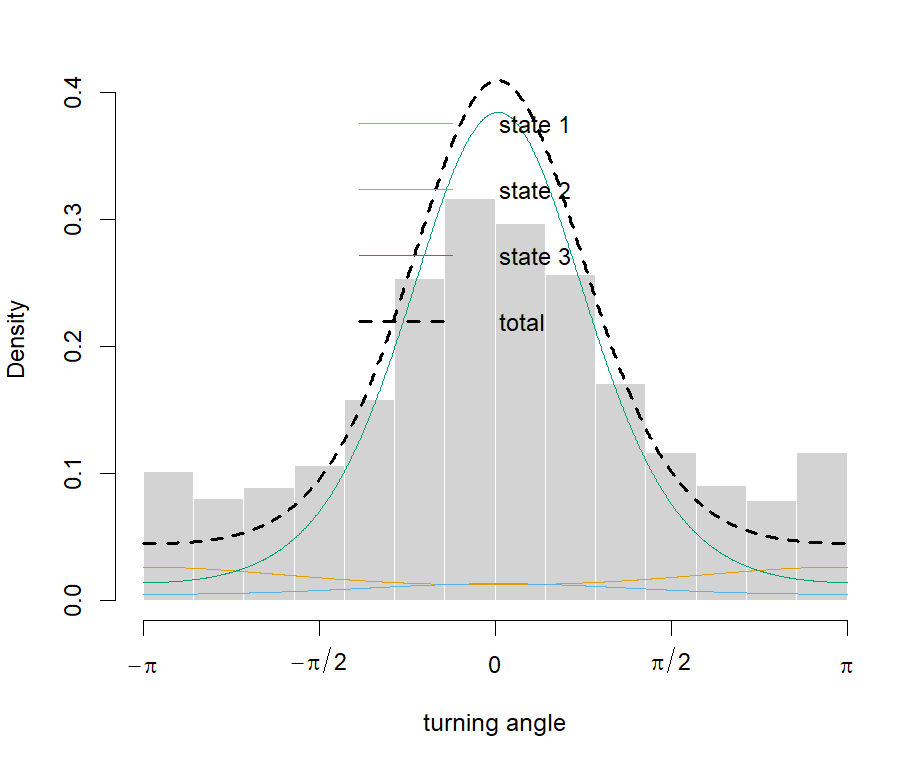

**b**

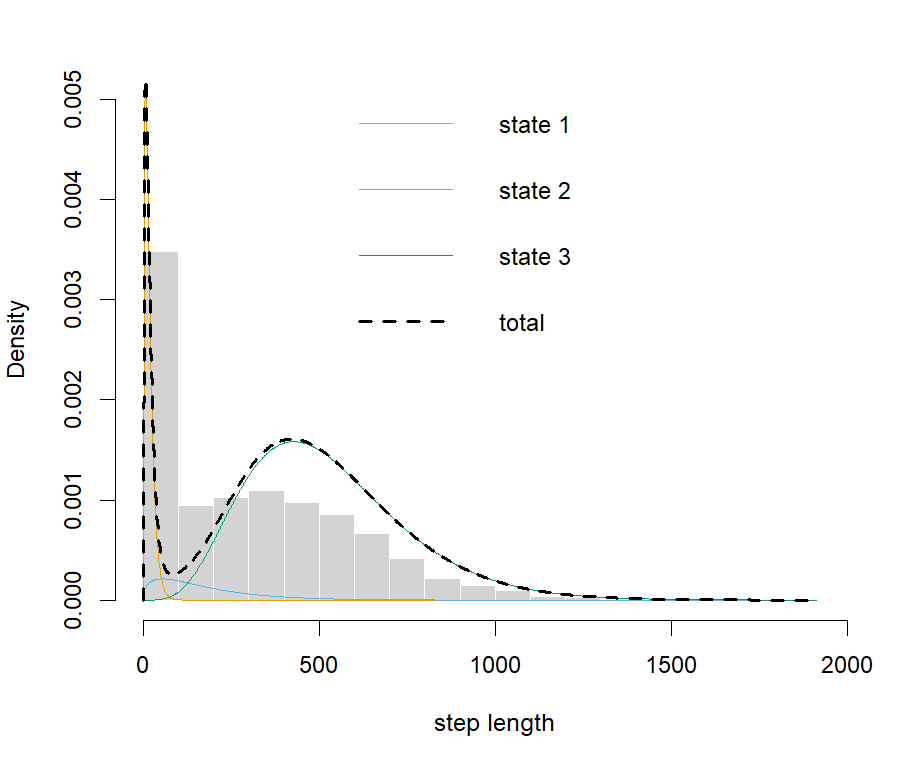

Appendix S4: Density of turning angles (a) and step lengths (b) for each of the 3 states in HMM models for female devils.

Appendix S5: Results of Manly’s habitat selection test for adult female devils for the 3 Hidden Markov Model behavioral states, as well as the proportion of availability for habitat types, and number and proportion of location fixes within each habitat type.

| Variable | State | Chi-squared test | df | p value | Category | Mean proportion of availability within MCPs | Number of location fixes | Proportion of location fixes (per variable/state) | Selection ratio | Lower 95% CI | Upper 95% CI |
| --- | --- | --- | --- | --- | --- | --- | --- | --- | --- | --- | --- |
| Distance to road | 1 | 36.79 | 20 | 0.01 | <20 m | 0.05 | 43 | 0.13 | 2.71 | 1.38 | 4.05 |
|  |  |  |  |  | 20-50 m | 0.07 | 34 | 0.10 | 1.46 | 0.83 | 2.10 |
|  |  |  |  |  | >50 m | 0.89 | 261 | 0.77 | 0.87 | 0.79 | 0.95 |
|  | 2 | 40.11 | 22 | 0.01 | <20 m | 0.05 | 54 | 0.12 | 2.48 | 1.59 | 3.37 |
|  |  |  |  |  | 20-50 m | 0.07 | 54 | 0.12 | 1.69 | 1.19 | 2.20 |
|  |  |  |  |  | >50 m | 0.89 | 361 | 0.77 | 0.87 | 0.81 | 0.92 |
|  | 3 | 62.88 | 22 | <0.001 | <20 m | 0.05 | 98 | 0.14 | 2.97 | 2.10 | 3.82 |
|  |  |  |  |  | 20-50 m | 0.07 | 50 | 0.07 | 1.03 | 0.57 | 1.49 |
|  |  |  |  |  | >50 m | 0.89 | 564 | 0.79 | 0.89 | 0.83 | 0.96 |
| Distance to plantation edge | 1 | 70.89 | 24 | <0.001 | <20 m | 0.13 | 114 | 0.34 | 2.38 | 1.66 | 3.10 |
|  |  |  |  |  | 20-50 m | 0.16 | 66 | 0.20 | 1.16 | 0.89 | 1.43 |
|  |  |  |  |  | >50 m | 0.71 | 158 | 0.47 | 0.68 | 0.47 | 0.89 |
|  | 2 | 90.11 | 24 | <0.001 | <20 m | 0.13 | 146 | 0.31 | 2.24 | 1.63 | 2.85 |
|  |  |  |  |  | 20-50 m | 0.16 | 113 | 0.24 | 1.47 | 1.11 | 1.82 |
|  |  |  |  |  | >50 m | 0.71 | 210 | 0.45 | 0.64 | 0.49 | 0.79 |
|  | 3 | 176.94 | 24 | <0.001 | <20 m | 0.13 | 278 | 0.39 | 2.78 | 2.09 | 3.46 |
|  |  |  |  |  | 20-50 m | 0.16 | 128 | 0.18 | 1.08 | 0.79 | 1.37 |
|  |  |  |  |  | >50 m | 0.71 | 306 | 0.43 | 0.62 | 0.49 | 0.75 |
| Habitat type | 1 | 21.63 | 22 | 0.48 | Plantation | 0.38 | 117 | 0.35 | 0.97 | 0.71 | 1.23 |
|  |  |  |  |  | Native forest | 0.46 | 131 | 0.39 | 1.06 | 0.84 | 1.28 |
|  |  |  |  |  | Grassland | 0.11 | 28 | 0.08 | 0.88 | 0.48 | 1.29 |
|  | 2 | 17.29 | 21 | 0.69 | Plantation | 0.38 | 165 | 0.35 | 1.00 | 0.77 | 1.22 |
|  |  |  |  |  | Native forest | 0.46 | 181 | 0.39 | 1.04 | 0.85 | 1.23 |
|  |  |  |  |  | Grassland | 0.11 | 39 | 0.08 | 0.85 | 0.58 | 1.12 |
|  | 3 | 22.27 | 24 | 0.56 | Plantation | 0.38 | 265 | 0.37 | 1.10 | 0.95 | 1.25 |
|  |  |  |  |  | Native forest | 0.46 | 216 | 0.30 | 0.87 | 0.71 | 1.03 |
|  |  |  |  |  | Grassland | 0.11 | 75 | 0.11 | 1.12 | 0.84 | 1.40 |
| Plantation ages | NA | 77.99 | 30 | <0.001 | <1 year | 0.01 | 19 | 0.05 | 1.08 | 0.87 | 1.29 |
|  |  |  |  |  | 1-3 years | 0.09 | 97 | 0.26 | 0.96 | 0.47 | 1.46 |
|  |  |  |  |  | 4-7 years | 0.08 | 149 | 0.40 | 1.52 | 0.98 | 2.05 |
|  |  |  |  |  | 8-13 years | 0.05 | 39 | 0.11 | 0.74 | 0.28 | 1.20 |
|  |  |  |  |  | 14+ years | 0.11 | 66 | 0.18 | 0.65 | 0.16 | 1.15 |

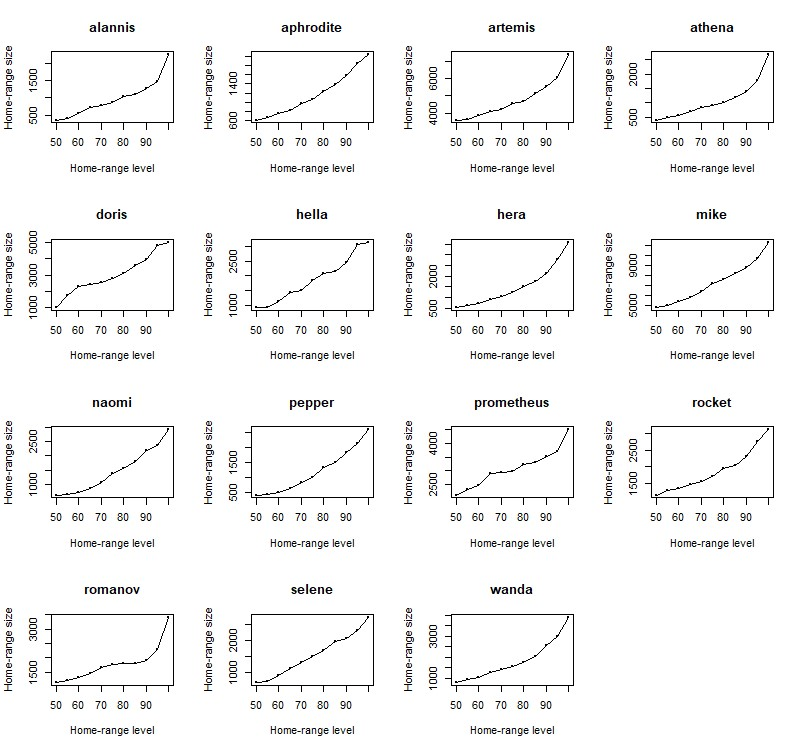

Appendix S6: Minimum convex polygon plots showing the increase in home range size (ha) vs % of retrieved location points included (home-range level).

Appendix S7: All final candidate models for predicting devil home range size and velocity in plantations, with square brackets representing 95% CIs**.**

| Home range size (ha): all devils | | | | | | | | | | | |
| --- | --- | --- | --- | --- | --- | --- | --- | --- | --- | --- | --- |
| Fixed effects | | | | | | | | | | Model description | |
| Intercept | | | | | Sex | | | | | df | AICc |
| 1849 [1433 to 2264] (female) | | | | | 1494 [566 to 2421] (male) | | | | | 3 | 234.6 |
| Home range size (ha): excluding male with largest home range | | | | | | | | | | | |
| Model | Fixed effects | | |  | | | | | | Model description | |
|  | Intercept | | | | Sex | | | | | df | AICc |
| #1 | 1849 [1585 to 2113] (female) | | | | 714 [16 to 1412] (male) | | | | | 3 | 212.7 |
| #2 (Null) | 1585 [1674 to 2228] | | | | |  | | | | 2 | 215.5 |
| Velocity (m.sec^-1^): whole-of-landscape | | | | | | | | | | | |
| Fixed effects | | | | | | | | Random effects | | Model description | |
| Intercept (m) | | Distance to road (m) | | Distance to plantation edge (m) | | | | Intercept (animal ID) | Residual | df | AIC |
| 0.41 [0.38 to 0.44] (< 20) | | -0.09 [-0.12 to  -0.06] (20–50)  -0.06 [-0.09 to  -0.04] (> 50) | | -0.07 [-0.09 to 0.05] (20–50)  -0.08 [-0.10 to  -0.06] (> 50) | | | | 0.002 (0.04 SD) | 0.09 (0.30 SD) | 5 | 2511.3 |
| Velocity (m.sec^-1^): plantation only | | | | | | | | | | | |
| Fixed effects | | | | | | | | Random effects | | Model description | |
| Intercept | | | Distance to plantation edge (m) | | | |  | Intercept (animal ID) | Residual | df | AICc |
| 0.38 [0.34 to 0.43] (< 20 m) | | | -0.10 [-0.13 to -0.06] (20-50)  -0.10 [-0.13 to -0.07] (> 50) | | | |  | 0.01 (0.08 SD) | 0.09 (0.30 SD) | 5 | 924.3 |
